# A large-scale whole-genome sequencing analysis reveals highly specific genome editing by both Cas9 and Cpf1 nucleases in rice

**DOI:** 10.1101/292086

**Authors:** Xu Tang, Guanqing Liu, Jianping Zhou, Qiurong Ren, Qi You, Li Tian, Xuhui Xin, Zhaohui Zhong, Binglin Liu, Xuelian Zheng, Dengwei Zhang, Aimee Malzahn, Zhiyun Gong, Yiping Qi, Tao Zhang, Yong Zhang

**Affiliations:** Department of Biotechnology, School of Life Sciences and Technology, Center for Informational Biology, University of Electronic Science and Technology of China, Chengdu 610054, China; Jiangsu Key Laboratory of Crop Genetics and Physiology, Co-Innovation Center for Modern Production Technology of Grain Crops, Key Laboratory of Plant Functional Genomics of the Ministry of Education, Yangzhou University, Yangzhou 225009, China; Joint International Research Laboratory of Agriculture and Agri-Product Safety of Ministry of Education of China, Yangzhou University, Yangzhou 225009, China; Department of Plant Science and Landscape Architecture, University of Maryland, College Park, Maryland 20742, USA; Institute for Bioscience and Biotechnology Research, University of Maryland, Rockville, Maryland 20850

**Author notes:** These authors contributed equally to this work (Xu Tang, Guanqing Liu, Jianping Zhou). Corresponding authors: Yiping Qi, Department of Plant Science and Landscape Architecture, University of Maryland, College Park, MD 20742, USA; Tao Zhang, Jiangsu Key Laboratory of Crop Genetics and Physiology, Co-Innovation Center for Modern Production Technology of Grain Crops, Key Laboratory of Plant Functional Genomics of the Ministry of Education, Yangzhou University, Yangzhou 225009, China; Yong Zhang, Department of Biotechnology, School of Life Sciences and Technology, Center for Informational Biology, University of Electronic Science and Technology of China, Room 216, Main Building, No. 4, Section 2, North Jianshe Road, Chengdu, 610054, P.R. China.

## Abstract

**Targeting specificity has been an essential issue for applying genome editing systems in functional genomics, precise medicine and plant breeding. Understanding the scope of off-target mutations in Cas9 or Cpf1-edited crops is critical for research and regulation. In plants, only limited studies had used whole-genome sequencing (WGS) to test off-target effects of Cas9. However, the cause of numerous discovered mutations is still controversial. Furthermore, WGS based off-target analysis of Cpf1 has not been reported in any higher organism to date. Here, we conducted a WGS analysis of 34 plants edited by Cas9 and 15 plants edited by Cpf1 in T0 and T1 generations along with 20 diverse control plants in rice, a major food crop with a genome size of ~380 Mb. The sequencing depth ranged from 45X to 105X with reads mapping rate above 96%. Our results clearly show that most mutations in edited plants were created by tissue culture process, which caused ~102 to 148 single nucleotide variations (SNVs) and ~32 to 83 insertions/deletions (indels) per plant. Among 12 Cas9 single guide RNAs (sgRNAs) and 3 Cpf1 CRISPR RNAs (crRNAs) assessed by WGS, only one Cas9 sgRNA resulted in off-target mutations in T0 lines at sites predicted by computer programs. Moreover, we cannot find evidence for bona fide off-target mutations due to continued expression of Cas9 or Cpf1 with guide RNAs in T1 generation. Taken together, our comprehensive and rigorous analysis of WGS big data across multiple sample types suggests both Cas9 and Cpf1 nucleases are very specific in generating targeted DNA modifications and off-targeting can be avoided by designing guide RNAs with high specificity**.

---

Bacterial type II CRISPR-Cas9 systems can effectively induce RNA-guided DNA double strand breaks (DSBs)^1^, making them popular tools for genome editing in bacteria^2^, animal cells^3^, mammalian systems^4–7^ and plants^8–11^. The most widely used *Streptococcus pyogenes* Cas9 (SpCas9) uses ~20 nucleotides (nt) of a single guide RNA (sgRNA) to recognize a complementary target DNA site along with an NGG protospacer adjacent motif (PAM)^1, 12^. More recently, type V CRISPR-Cpf1 was shown to mediate efficient genome editing in human cells^13^ and plants^14, 15^. Cpf1 uses ~23 nt of an RNA guide to target DNA with a TTTV PAM^13^. RNA-guided nucleases (RGNs) such as Cas9 and Cpf1 represent versatile genome editing tools that promise to advance basic science, enable personalized medicine and accelerate crop breeding. However, Cas9 may cause undesired off-target mutations due to sgRNAs recognizing DNA sequences with one to a few nucleotide mismatches; albeit with reduced nuclease binding and cleavage activity^1, 6, 16, 17^. Although similar rules apply to Cpf1, recent studies in human cells^18, 19^ have shown Cpf1 is generally more specific than Cas9.

Understanding the scope of off-target mutations in Cas9 or Cpf1-edited crops is critical for research and regulation. Previously, whole-genome sequencing (WGS) was applied for detecting off-target mutations by Cas9 in *Arabidopsis*^20^, rice^21^ and tomato^22^. Unfortunately, these studies either only looked at potential off-target sites predicted by computer programs or fell short of full analysis of all the mutations identified by WGS in edited plants. Without inclusion of enough necessary controls, such WGS studies had limited power for isolating off-target mutations in edited plants because they were unable to fully assess the levels of preexisting mutations, spontaneous mutations, and mutations caused by tissue culture and *Agrobacterium* mediated transformation. Genome-wide identification of off-target mutations by Cas9 or Cpf1 will be empowered only if all these background mutations can be isolated. Furthermore, WGS based off-target analysis of Cpf1 has not been reported in any higher organism. In recent years, WGS studies on Cas9-edited mice have generated contrasting results; one study found few off-target mutations^23^ while the other found many^24^. This controversy raised the urgency for comprehensive and rigorous analyses of off-target mutations using WGS in edited animals and plants. We reasoned a large-scale and well-designed study is required for comprehensive assessment and comparison of off-target effects by Cas9 and Cpf1 in crops. Here, we describe a large-scale WGS study to assess off-target effects of Cas9 and Cpf1 in rice, an important food crop. Our results suggest off-target effects of Cas9 and Cpf1 are largely negligible when compared to spontaneous mutations or mutations caused by tissue culture and *Agrobacterium* infection in edited plants. The resulting knowledge is likely to serve as an important reference for plant researchers and regulatory agencies.

## RESULTS

### Detection of off-target, spontaneous and background mutations

To comprehensively evaluate potential off-target effects of Cas9 in rice, we generated 10 T-DNA constructs to target 7 genes with 12 sgRNAs including two dual-sgRNA constructs for editing two circular RNA loci (**Supplementary Table 1**). All 10 CRISPR-Cas9 nuclease expression constructs were active at target sites and resulted in editing frequencies ranging from 15% to 100% in T0 lines (**Fig. 1a, 1b and Supplementary Table 1**). For each Cas9 construct, two independent T0 plants carrying non-mosaic mutations (**Supplementary Fig. 1**) were chosen for WGS. To assess off-target effects of Cpf1, we followed three previously published *Lachnospiraceae bacterium* ND2006 Cpf1 (LbCpf1) targeting constructs that resulted in 100% editing efficiency in T0 lines^14^. Two Cpf1 T0 plants per construct carrying non-mosaic on-target mutations were chosen for WGS (**Supplementary Fig. 2**). For four T0 lines edited by four different Cas9 sgRNAs and two T0 lines edited by two different Cpf1 crRNAs, we selected two to five plants from each T0 line in the T1 generation for WGS (**Fig. 1a and 1b**). In addition, four wild type (WT) plants each from three consecutive generations were also included for WGS to survey spontaneous mutations (**Fig. 1b**). To ensure high confidence on base calling, all 69 individual plants were sequenced at 45X to 105X in depth (**Supplementary Table. 2**). A stringent mutation mapping and calling pipeline was developed for WGS analysis (**Fig. 1c**). Single-nucleotide variants (SNVs) and small insertions and deletions (indels) were each identified with three variant-calling software programs, with high-confident variants shared by all software being further analyzed for mutation identification (**Fig. 1c**).

**Figure 1.**
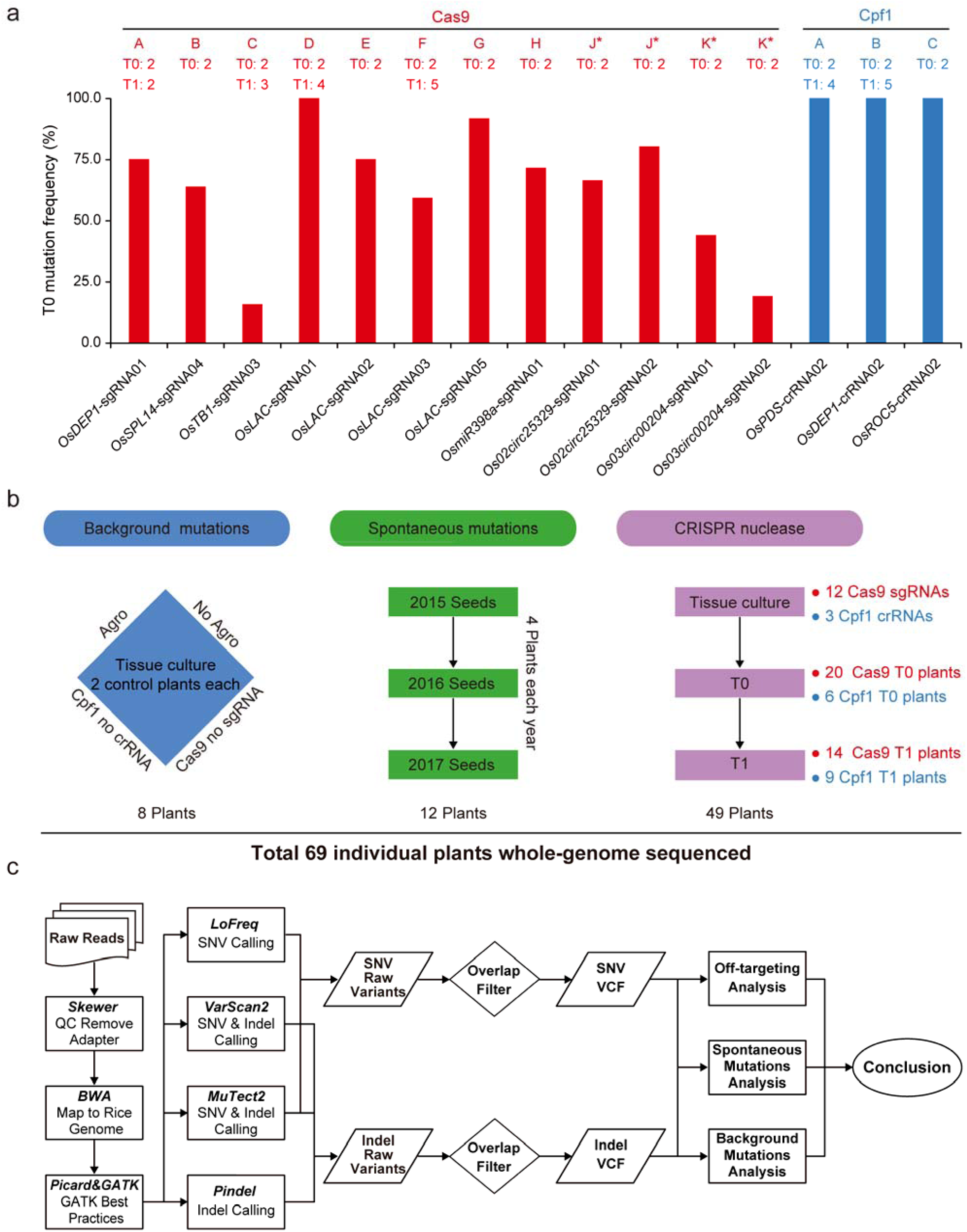
Experimental design and work flow. (**a**) Genome editing efficiency at selected 12 Cas9 and 3 Cpf1 target sites in T0 rice plants. The X-axis shows the names of sgRNAs and crRNAs which are denoted as Cas9-A to Cas9-K and Cpf1-A to Cpf1-C. The numbers of T0 and/or T1 lines that are subjected to whole-genome sequencing (WGS) are indicated. The Y-axis shows genome editing frequencies calculated based on genotyping data in T0 generation. *Cas9-J and Cas9-K samples each express a dual-sgRNA construct, targeting two genes simultaneously. (**b**) Selection of plants for WGS. Left: four groups of controls are included for assessing different background mutations. Middle: three generations of wild type plants are included for assessing parent-progeny spontaneous mutations. Right: Multiple T0 and T1 lines edited by Cas9 and Cpf1 are chosen for assessing off-targeting by WGS. (**c**) Workflow of whole-genome detection of SNV and indel mutations. SNV analysis involves using three computer programs: LoFreq, VarScan2 and MuTect2. Indel analysis also involves using three programs: VarScan2, MuTect2 and Pindel.

**Figure 2.**
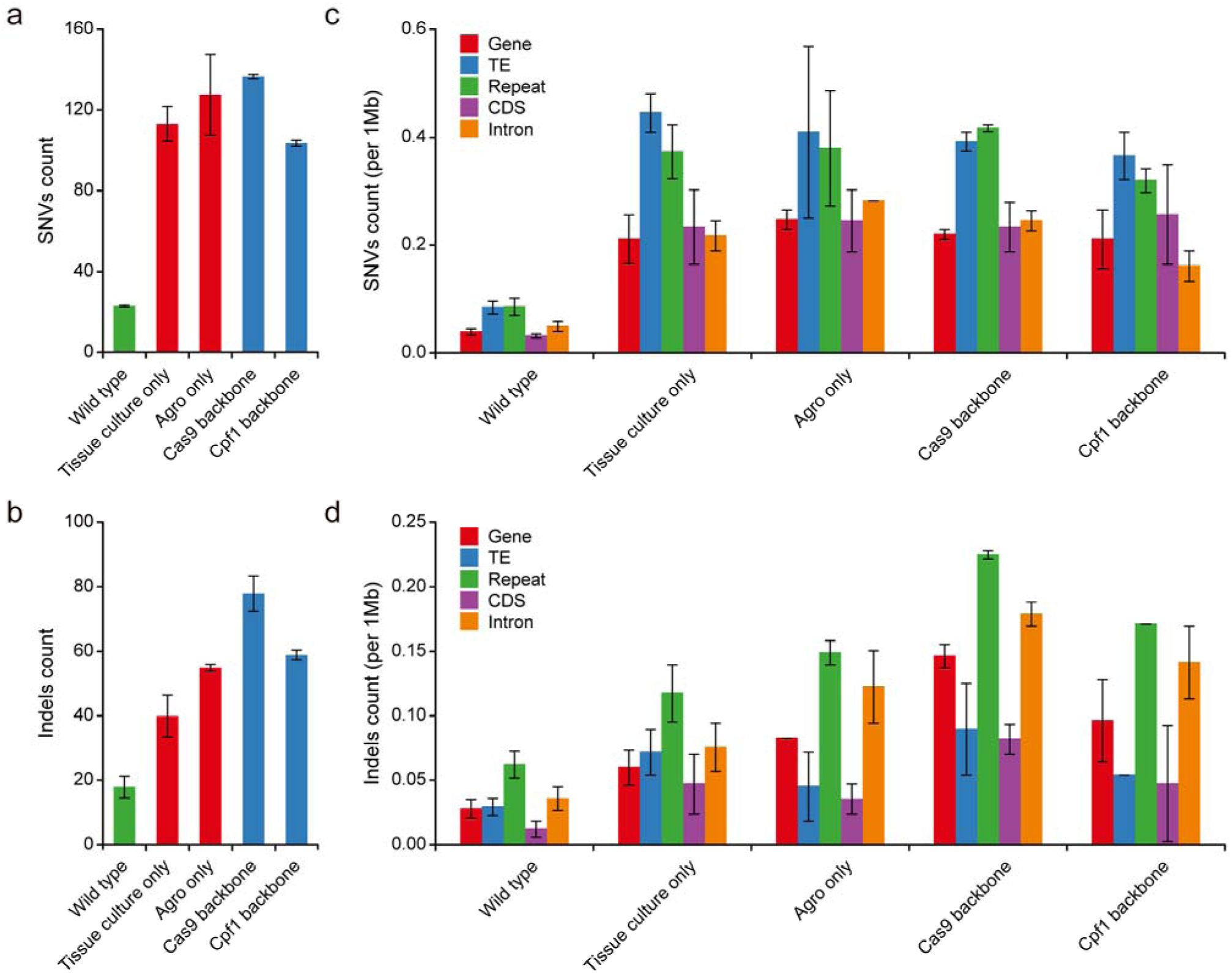
Genome-wide analysis of spontaneous mutations and mutations caused by tissue culture and Agrobacterium mediated transformation. (**a, b**) Average numbers of SNVs and indels detected in 3 generations of wild type plants and 4 types of tissue culture-related control plants. Error bars indicate s.e.m. (**c, d**) Annotation of genome-wide distribution of mutations found in all control samples: WT, tissue culture only, Agro-infection, Cas9 backbone and Cpf1 backbone. TE, transposable element. CDS, coding sequence. Error bars indicate s.e.m.

To survey pre-existing mutations in the WT population and estimate the level of spontaneous mutations across generations, we analyzed the WGS data from 12 WT plants across three consecutive generations (**Fig. 1b and Supplementary Fig. 3**) After filtering shared pre-existing mutations, we estimated an average of 23 SNVs and 18 indels as spontaneous mutations from parents to progeny in rice (**Fig. 2a and 2b**). We calculated the spontaneous mutation rate at ~5.4×10^−8^ per site per diploid genome per generation, which is in line with the rates previously reported in maize (2.2-3.9×10^−8^)^25^ but higher than the rate in *Arabidopsis* (7-7.4×10^−9^)^26, 27^.

**Figure 3.**
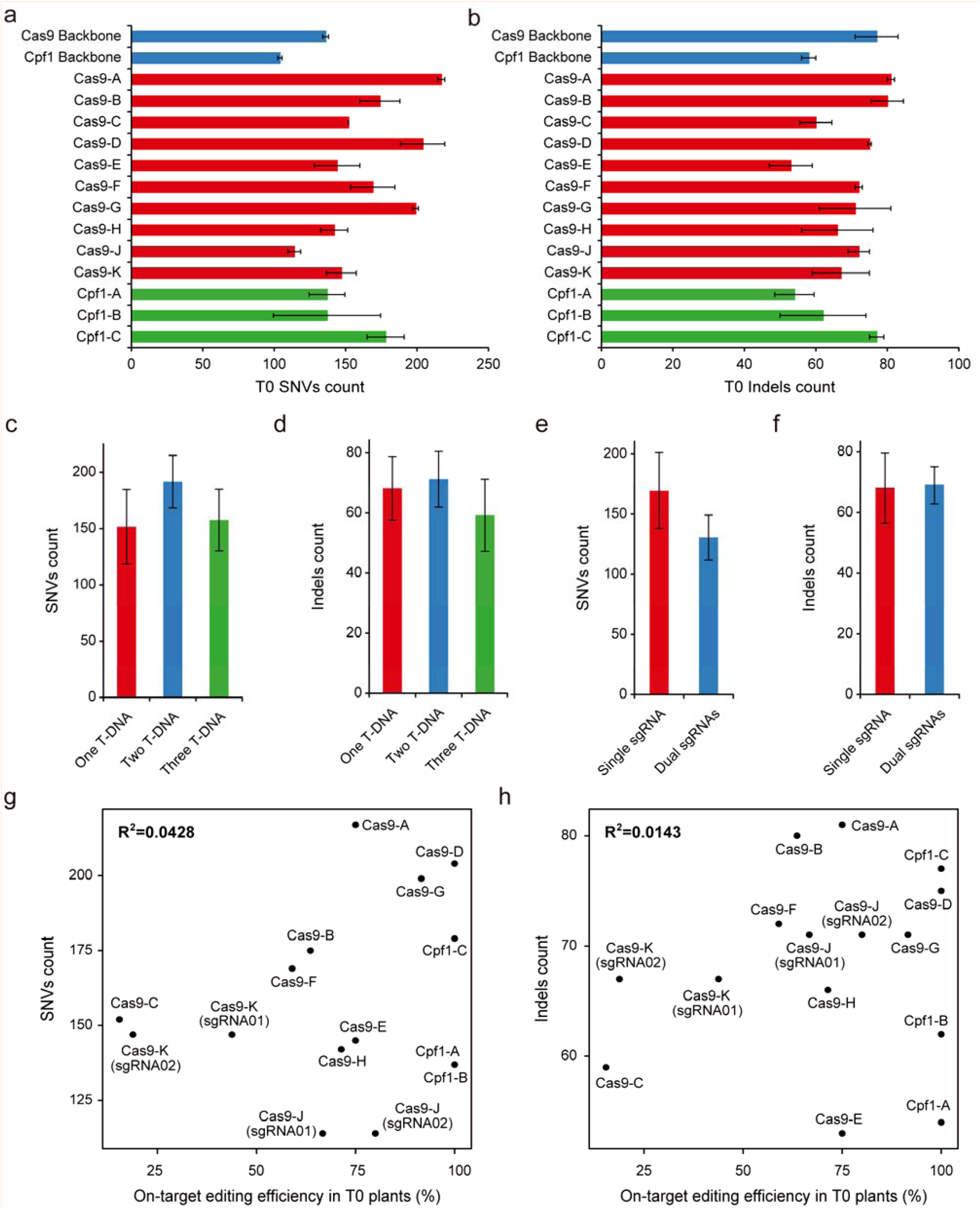
Detailed analysis of mutations at Cas9 or Cpf1-edited T0 plants. (**a**, **b**) Average numbers of SNVs and indels detected in 26 T0 plants edited by Cas9 or Cpf1. (**c**, **d**) Average numbers of SNVs and indels in edited T0 plants with different numbers of T-DNA insertions. (**e**, **f**) Average numbers of SNVs and indels in Cas9-edited T0 plants expressing one or two sgRNAs (in Cas9-J and Cas9-K). (**g**, **h**) Pearson correlation between on-target editing frequency and the numbers of SNVs or indels mutations in Cas9 and Cpf1-edited T0 plants. Error bars in **a-f** indicate s.e.m.

To assess mutations generated by tissue culture and *Agrobacterium* infection, we produced and sequenced four types of control plants: tissue culture only, tissue culture with *Agrobacterium*, tissue culture with *Agrobacterium* transformation of Cas9 without sgRNA, and tissue culture with *Agrobacterium* transformation of Cpf1 without crRNA (**Fig. 1b and Supplementary Fig. 4**). Tissue culture is known to be mutagenic, causing somaclonal variations^28^. Indeed, the two tissue culture-only samples contained an average of 114 SNVs and 36 indels (**Fig. 2a** and **2b**), resulting in a background mutation rate of 1.86×10^−7^, which is similar to the rates (1.7-3.3×10^−7^) previously published^29^. Importantly, similar numbers of SNVs were observed from *Agrobacterium*-infected or Cas9/Cpf1 backbone-transformed plants (**Fig. 2a**). These three controls generated ~15 to 41 more indels compared to tissue culture-only samples (**Fig. 2b**), suggesting *Agrobacterium* infection is mutagenic with a preference for introducing indels. This warrants further investigation as these three controls show large variations on indel counts. We mapped all identified mutations from these four control types to the rice genome across 12 chromosomes (**Supplementary Fig. 5**). Further analysis of the genome-wide distribution of these background mutations revealed high enrichment of SNVs in transposable elements (TE) and repeats (**Fig. 2c**), as well as high enrichment of indels in repeats (**Fig. 2d**).

**Figure 4.**
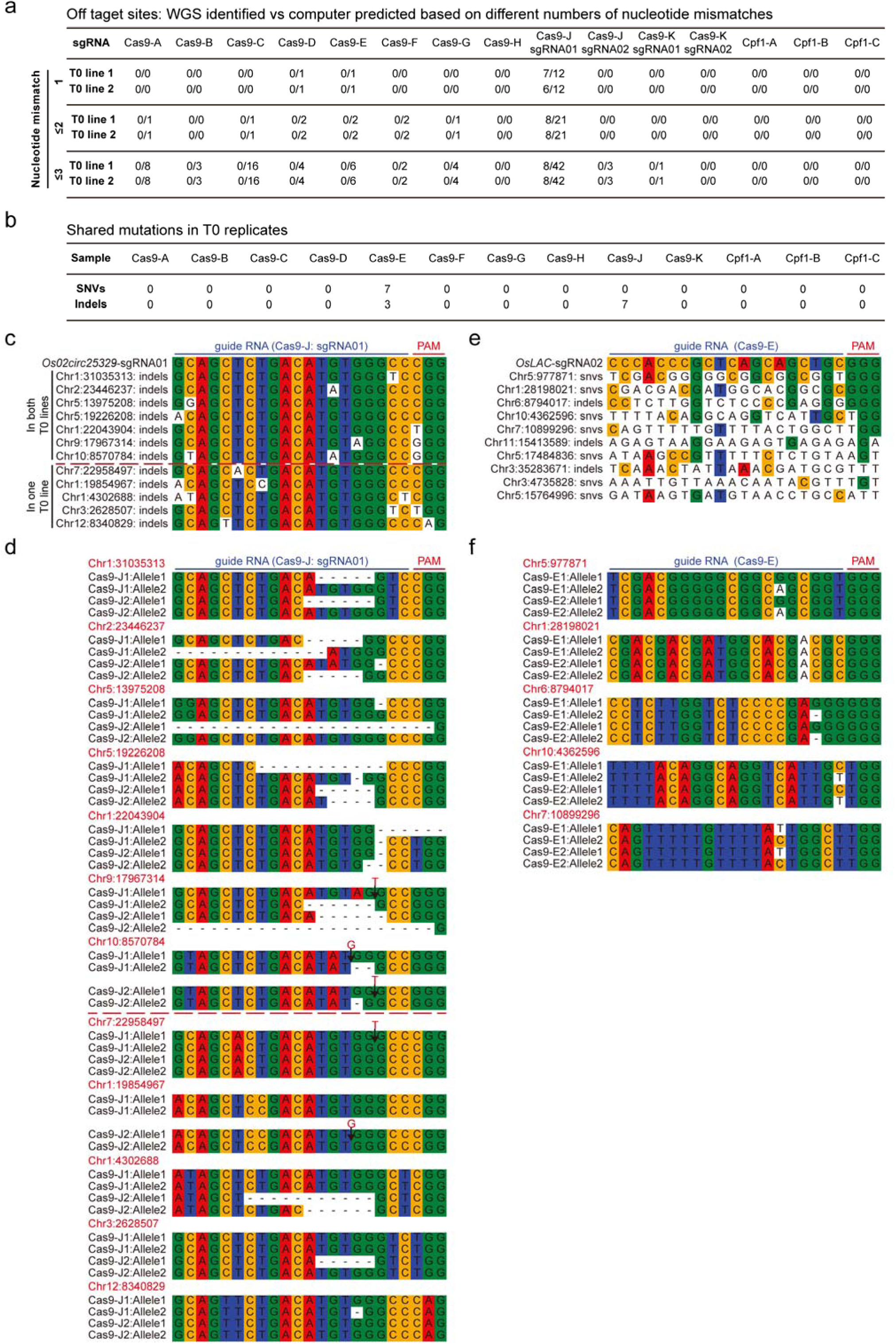
Analysis and identification of potential off-target sites in T0 lines. (**a**) Number of off-target sites identified in replicated T0 plants vs the number of all off-target sites that are predicted by Cas-OFFinder and CRISPOR with allowing up to 3 nt mismatch for all 15 Cas9 or Cpf1 target sites. (**b**) Identification of shared SNVs and indels between replicated T0 plants. (**c**) Potential off-target sites identified in both Cas9-J T0 samples (above the red dashed line) and only in one T0 sample (below the red dashed line). (**d**) Off-target mutations identified by WGS at off-target sites in both Cas9-J T0 samples (above the red dashed line) and only in one T0 sample (below the red dashed line). (**e**) Potential off-target sites identified in Cas9-E samples based on shared mutations in two T0 plants. (**f**) Sequence analysis of the shared mutations in Cas9-E samples.

**Figure 5.**
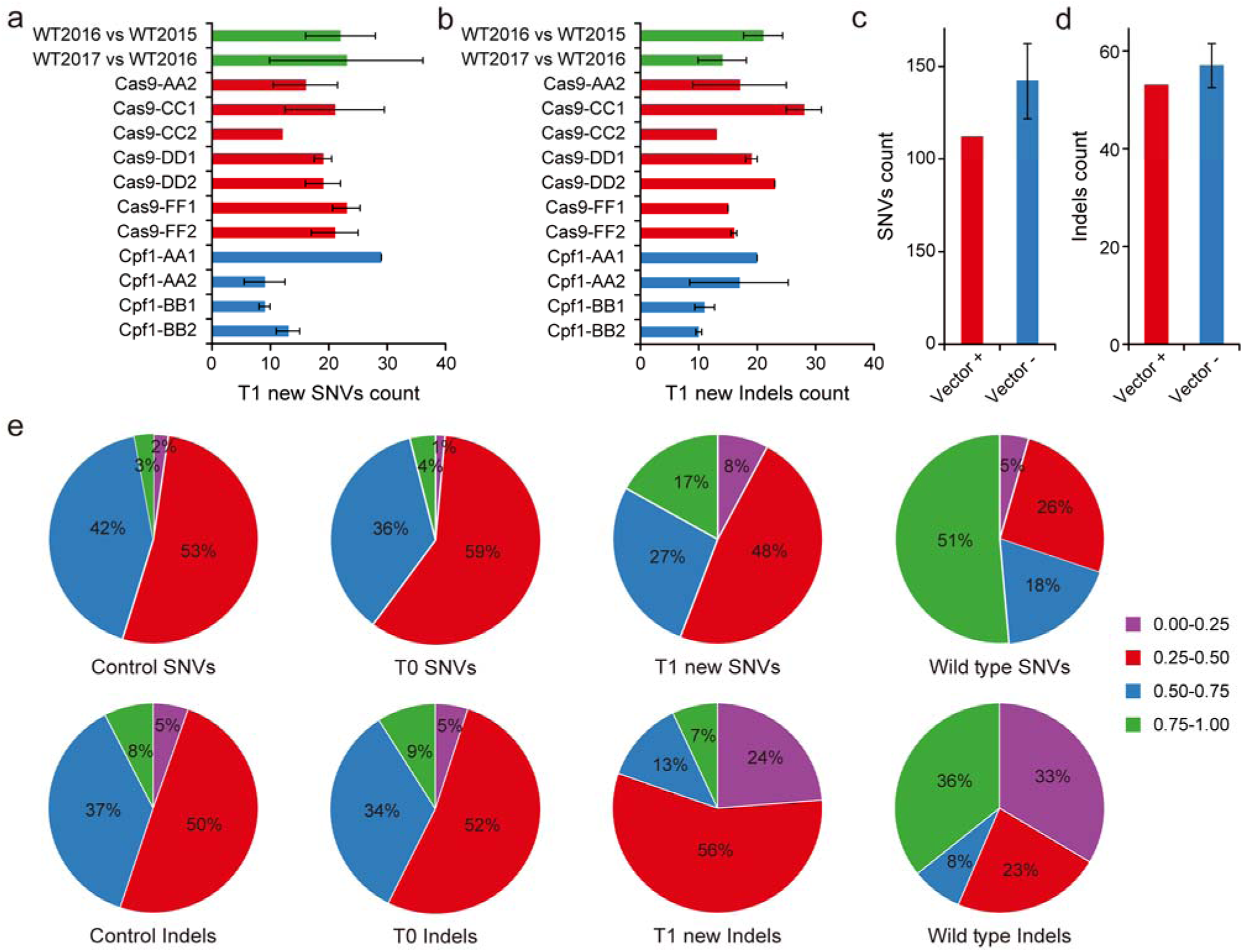
Evaluate off-target effect of Cas9 and Cpf1 in T1 plants. (**a,b**) Analysis of new SNVs and indels in T1 plants. (**c, d**) Analysis of SNVs and indels in T1 plants that carry (+) or do not carry (-) the Cas9 and sgRNA expression cassettes. (**e**) Allele frequencies of SNVs and indels identified in all tissue culture related controls (tissue culture and transformants with Cas9 and Cpf1 backbones), T0, T1 and WT plants. Above: SNVs; Bottom: indels.

### SNVs and indels identified in edited T0 plants are largely background mutations

Whole-genome sequencing of 20 Cas9 and 6 Cpf1-edited T0 lines confirmed all target site mutations that were initially identified with Sanger sequencing (**Supplementary Table 1, Supplementary Fig. 1 and 2**). We identified SNVs and indels in these Cas9 T0 lines (**Supplementary Fig. 6**) and Cpf1 T0 lines (**Supplementary Fig. 7**) and mapped these mutations to the rice genome (**Supplementary Fig. 5**). We found their numbers are close to those in Cas9 or Cpf1 backbone controls, with about twice as many SNVs as indels (**Fig. 3a** and **3b**). This mutation pattern is not consistent with Cas9 or Cpf1-generated mutations in rice which are largely indels^9, 14^. For example, all target site mutations in these selected 26 T0 lines are indels (**Supplementary Table 1, Supplementary Fig. 1 and 2**). The SNV and indel mutations in Cas9 and Cpf1-edited T0 samples share similar genome-wide distribution with the tissue culture related controls (**Supplementary Fig. 5**). We identified a total of 31 T-DNA insertion events in 26 T0 lines and found T-DNA copy numbers ranging from 1 to 3; most T0 lines had only one T-DNA insertion (**Supplementary Fig. 8**). No significant difference was found for the numbers of SNVs and indels among T0 lines with different T-DNA copy numbers (**Fig. 3c** and **3d**). Cas9-J and Cas9-K T0 lines each expressed a dual-sgRNA construct for simultaneous expression of two sgRNAs, targeting two putative circle RNA genes (**Fig. 1a**). No significant difference was found for the numbers of SNVs and indels in these four dual-sgRNA lines and the other 22 single sgRNA lines (**Fig. 3e** and **3f**). Moreover, there is no correlation between the numbers of SNVs or indels and the on-target editing efficiency by Cas9 or Cpf1 in these T0 plants (**Fig. 3g** and **3h**). All these analyses strongly suggest mutations in these genome-edited T0 lines are mostly background mutations caused during tissue culture and *Agrobacterium* mediated transformation.

### Identification of true off-target mutations in T0 plants

To identify true off-target mutations in the T0 plants, we first evaluated the specificity of 12 sgRNAs of Cas9 and 3 crRNAs of Cpf1 with CRISPOR^30^ and Cas-OFFinder^31^. With a stringent criterion allowing only a 1 nt mismatch in the protospacer, three Cas9 sgRNAs (Cas9-D, Cas9-E and Cas9-J-sgRNA01) (**Fig. 1a**) had predicted off-target sites (**Fig. 4a** and **Supplementary Table 3**). When we mapped all identified mutations to these potential off-target sites by allowing up to 10 nt mismatches to the protospacers of Cas9 (**Supplementary Fig. 9**) and Cpf1 (**Supplementary Fig. 10**), only Cas9-J-sgRNA01 showed evidence of true off-targeting. It is worth noting that these off-target sites showed high sequence homology to the Cas9-J-sgRNA01 target site and could be accurately predicted by software such as CRISPOR and Cas-OFFinder (Supplementary **Table 3**). We reasoned true off-target mutations are likely to occur separately in independent T0 lines. Indeed, among 12 off-target sites identified for Cas9-J-sgRNA01, seven sites were overlapped between two T0 lines while the remaining five sites were only validated from one T0 line (**Fig. 4b and 4c**). All 12 off-target sites show very high sequence homology with the target site (**Fig. 4c**). Among them, one site at Chr1:22043904 is technically an on-target site because it has the same 20-nt protospacer with 1-nt silent mismatch in the PAM (CGG vs TGG). For the remaining 11 true off-target sites, eight sites carry one mismatch mutation in the 20 nt protospacer. For the additional 3 sites with two or three mismatch mutations, only one mutation is present in the 1-18 nt sequence from the PAM (**Fig. 4c**). Further analysis these 12 off-target sites found four have silent mutations in NGG PAM and one has a non-canonical CAG PAM, which was reported as an alternative PAM (NAG) for SpCas9 nuclease^32^ and recently shown to mediate Cas9 activity in rice^33^. All mutations at these 12 sites were indels, and, importantly, the two Cas9-J T0 lines carried distinct alleles at these sites (**Fig. 4d and Supplementary Fig. 11**); validating these mutations were truly caused by Cas9.

Cas9-E sgRNA was predicted by CRISPOR and Cas-OFFinder to contain 6 off-target sites when up to a 3 nt mismatch was allowed (**Fig. 4a and Supplementary Table 3**). However, no off-target mutations were found at these predicted sites. Although the two Cas9-E T0 lines shared seven SNVs and three indels (**Fig. 4b**), these 10 shared mutations had very poor sequence homology to the target site (**Fig. 4e**). Only five sites contained the NGG PAM. Among them, the site sharing highest sequence homology with the target site still contained a 10 nt mismatch, making it unlikely to be a true off-target site. Unlike indels found in Cas9-J samples, these putative off-target mutations are mostly SNVs (**Fig. 4b and f**). Furthermore, both independent T0 lines always carried the same mutant alleles (**Fig. 4f and Supplementary Fig. 11**). These observations suggest that the 10 shared mutations of two Cas9-E T0 lines were not caused by Cas9, but were pre-existing mutations from a parental line.

Cas9 was previously shown to induce off-target mutations at sites with missing or extra nucleotides when compared to the target site, which form bulges when targeted by guide RNAs^34^. To detect such off-target mutations, we extracted all T0 mutation site flanking sequences (25 bp upstream and downstream) and aligned them to corresponding sgRNA/crRNA sequences using BLAST. Only Cas9-J1 and Cas9-J2 samples had alignments to the Cas9-J-sgRNA01 target (15 in Cas9-J1 and 10 in Cas9-J2); other samples had no hit. None of the detected mutations were caused by bulge-forming DNA-sgRNA recognition. We also investigated whether DNA translocation events were induced by Cas9 or Cpf1 by searching for structural variants (SVs) and gene fusion events in the whole rice genome. We did not detect any translocation event in all T0 lines. Given the level of nuclease-induced DNA translation can be used for assessing targeting specificity^35^, absence of detectable translation events in all T0 samples here indicates these Cas9 and Cpf1 reagents are indeed very specific; limiting cleavage activity almost exclusively to the target sites.

### No evidence of off-target mutations in T1 plants

Our analysis of T0 plants suggested 11 out of 12 Cas9 sgRNAs and all three Cpf1 crRNAs are very specific as no off-target mutations were detected. However, lack of off-target mutations might be attributed to low expression or activity of Cas9 or Cpf1. It is also important to determine whether continued expression of the RGNs into the next generation will result in *de novo* off-target mutations. Therefore, we decided to sequence 14 T1 plants from Cas9 T0 lines with diverse levels of on-target editing efficiency (15%, 60%, 75% and 100%) at four target sites and 9 T1 plants from Cpf1 T0 lines at two target sites (**Fig. 1a, 1b** and **Supplementary Fig. 1**). Germline-transmitted on-target mutations in 14 Cas9 edited or 9 Cpf1 edited T1 lines were validated by Sanger sequencing (**Supplementary Fig. 12 and 13**). With WGS analysis, we identified all SNVs and indels in Cas9 T1 lines (**Supplementary Fig. 14**) and Cpf1 T1 lines (**Supplementary Fig. 15**). The GWS results confirmed the germline-transmitted on-target mutations (**Supplementary Table 1, Supplementary Fig. 12 and 13**). Among all other SNVs and indels, most of them were identified in the corresponding T0 lines, suggesting they have been fixed (**Supplementary Fig. 16**). For the other new mutations identified in T1 lines, the average number of SNVs ranged from 9 to 29 (**Fig. 5a**), while the average of indels ranged from 10 to 28 (**Fig. 5b**). Such spontaneous mutation rates are consistent with the spontaneous mutation rates we found earlier in WT samples (**Fig. 2a and 2b**), which are also in line with a previous study^27^.

These new mutations were mapped to the rice genome alongside with new mutations that were discovered in WT plants across two generations (**Supplementary Fig. 17**). The genome distribution of these new mutations in T1 lines also showed enrichment in repeats (**Supplementary Fig. 16**), consistent with the spontaneous mutations discovered in WT (**Fig. 2c and 2d**). Detailed analysis of SNVs among all sample types revealed T1 lines have higher rates of G:C>A:T transitions than T0 lines (**Supplementary Fig. 18**), consistent with the observation on spontaneous mutations in *Arabidopsis*^26^. Further analysis of T1 lines either with or without the Cas9 transgene did not reveal any difference on the numbers of new SNVs and indels among these two subpopulations (**Fig. 5c and d**). By applying similar methods from the analysis of T0 plants, we were unable to identify any off-target mutations by Cas9 or Cpf1 in T1 lines. Given most T1 lines analyzed still carry the RGN constructs, our results suggest continued expression of Cas9 or Cpf1 constructs did not cause *de novo* off-target mutations in T1 lines.

To further assess the new mutations found in T1 lines, we calculated and compared the allele frequency of SNVs and indels among four groups: tissue culture controls, T0 plants, T1 plants, and WT (**Fig. 5e**). The tissue culture controls and Cas9/Cpf1 T0 lines share strikingly similar (mostly heterozygous-like) allele frequency distribution. This reiterates our earlier conclusion that all mutations in T0 samples (except a few found in Cas9-J samples) are background mutations. By contrast, T1 plants show more homozygous-like SNVs (0.75 to 1.0 in allele frequency) and somatic-like indels (0 to 0.25 in allele frequency). This trend of rapidly fixing SNVs and the increase of somatic indels in T1 is interesting, and relatively in line with the observation in WT plants.

## Discussion

Specificity of CRISPR-Cas RGN systems has caught more attentions in humans than in animals or plants, due to medicinal applications of RGNs. Earlier WGS studies in human cells found low incidence of off-target mutations by Cas9^36, 37^. Recently, two WGS off-target studies in mice showed conflicting results^23, 24^. However, the study that claimed unexpected large-scale off-target effects by Cas9 may be flawed due to limitations in its experimental design and WGS data analysis^24^. Given the wide adoption of CRISPR-Cas systems in agriculture, with genome-edited crop products reaching market in record time^38^, it becomes urgent to conduct large-scale and exhaustive WGS analysis of off-target effects by Cas9 and Cpf1, two leading RGN systems, in agriculturally important crops. Such studies will help assess safety of Cas9 and Cpf1 in precise crop breeding as well as provide valuable information to scientists, breeders, regulators and consumers.

In this study, we conducted a large-scale WGS analysis for detecting potential off-target mutations caused by 12 Cas9 sgRNAs and 3 Cpf1 crRNAs in rice, an important food crop. We confirmed WGS-identified mutations by Sanger sequencing at randomly selected sites with a 100% success rate (**Supplementary Table 4**), which is consistent with the high quality of our WGS data. Our experimental design took into account background mutations caused by tissue culture and *Agrobacterium* mediated transformation, pre-existing mutations in parents and spontaneous mutations that arise from seed propagation. Through sequencing 20 control plants of different types and 49 Cas9 or Cpf1-edited T0 and T1 plants, we only found true off-target mutations in two T0 lines expressing Cas9 protein with Cas9-J-sgRNA01. Importantly, these empirically validated off-target sites can be readily predicted computationally. Our examination of T1 plants that continue to carry Cas9-sgRNA or Cpf1-crRNA did not reveal off-target mutations, suggesting continued presence of the RGN reagents with varying activity in plants does not cause off-target mutations if the guide RNAs are well-designed for specificity. This observation is also highly significant because it encourages the use of Cas9 and Cpf1 in certain breeding applications that may require expression of RGNs across several generations. For example, a RGN cassette may be introduced from a transgenic line into a transformation-recalcitrant variety of the same plant species for genome editing with simple genetic crossing.

Our study also provided insights on avoiding off-target effects of Cas9 and Cpf1 in edited crops. To minimize off-target effects, many systems have been developed including paired Cas9 nickases^39^, high fidelity Cas9 proteins^40–42^, FokI-dCas9 fusions^43, 44^, truncated sgRNAs^45^, and ribonucleotide protein (RNP) delivery^46^. To assess and identify off-target sites, *in vivo*^17, 35^ and *in vitro*^47–49^ tools have also been developed in human cells, which may be applied in plants. Our WGS analysis with wild type SpCas9 and LbCpf1 proteins did not find off-target mutations for 14 out of 15 guide RNAs tested in T0 and T1 plants, suggesting utilization of a high-fidelity enzyme, which are typically of lower activity, may be unnecessary in crop applications. When a mismatch up to 3 nt of the protospacer is allowed, Cas9-OFFinder programs predicted a total of 37 off-target sites for 7 out of 11 Cas9 sgRNAs. Yet, we couldn’t detect any mutations at these putative off-target sites. Alternatively, Cas9-OFFinder predicted all the off-target sites that we identified for Cas9-J-sgRNA01; many of the sites have just 1 nt mismatch to the protospacer of the target site. Therefore, we can deduce a simple rule to alleviate off-target effects: making sure even the highest scored potential off-target sites will have at least a 2 nt mismatch to the seed sequence of the protospacer. We note this may not always be possible if the target sequence shares many homologous sequences in the genome. For example, maize has a very repetitive genome and wheat has A, B, D sub-genomes that share high similarity. In these cases, targeted amplicon sequencing using next generation sequencing technologies may be an appropriate and cost-effective method to look for off-target mutations.

Finally, we hope our data can be a valuable reference for regulatory agencies and other entities. It is reasonable and necessary to scrutinize any new technology for its efficacy and safety. Cas9 and Cpf1, as new crop breeding technologies, are no exception. Although Cas9 based off-target effects have been studied by WGS in plants^20–22^. our study differs from previous studies significantly at scale, depth and comprehensiveness. Our research also represents the first report of using WGS to assess off-targeting by Cpf1 in any edited higher eukaryotic organism. We could not find any off-target mutations in 47 out of 49 rice plants edited by 11 Cas9-sgRNA and 3 Cpf1-crRNA constructs. This precise level of genome modification casts a stunning contrast to many conventional breeding technologies. For example, we found that even the safest breeding approach, harvesting seeds from parental lines, introduces ~30 to 50 spontaneous mutations into the next generation in rice. We also observed ~200 tissue culture-introduced somaclonal variations per rice plant, even though few are affecting coding sequences. In conclusion, our data support a recent call to “Regulate genome-edited products, not genome editing itself”^50^.

## METHODS

### Plant Material and Growth Condition

This study used the rice variety Nipponbare (*Oryza sativa* L. ssp. Japonica cv. Nipponbare). All plants were grown in growth chambers under controlled environmental conditions with a 16/8 h light/dark regime at 28℃ and 60% relative humidity.

### Vector construction

Plasmids encoding for Cas9 and a single sgRNA were generated by ligating annealed oligos with a 4 bp overhang into a *Bsa*I digested backbone (either pZHY988 or pTX172)^51,52^. Plasmids with two sgRNAs were created by ligating pZHY988 with a 485 bp fragment, after digestion with *Bsa*I. This 485 bp fragment contains two sgRNAs generated by overlap extension PCR^52^. All CRISPR-Cpf1 nuclease expression vectors were reported in our previous study^14^. The sequences of all primers used to construct vectors are shown in **Supplementary Table 5**.

### Rice stable transformation

*Agrobacterium*-mediated rice transformation was performed as described in published protocols^53^ with slight modification. The binary vectors were introduced into *Agrobacterium tumefaciens* strain EHA105 by the freeze-thaw method^54^. For rice transformation, dehusked seeds were sterilized with 70% ethanol for 1 min. Afterwards, seeds were washed five times with sterile water, then further sterilized for 15 minutes with a 2.5% sodium hypochlorite solution containing a drop of Tween 20. The washing and sterilization step were repeated, this time without addition of Tween. Seeds were then rinsed an additional five times before being dried on sterilized filter paper and cultured on solid medium at 28°C in a dark growth chamber for 2-3 weeks. Actively growing calli were collected for subculture at 28°C in the dark for 1-2 weeks. *Agrobacterium* cultures were collected and resuspended in liquid medium (OD600=0.06-0.1) containing 100 μM acetosyringone. Rice calli were immersed in the *Agrobacterium* suspension for 30 min, then dried on sterilized filter paper and co-cultured for three days on solid medium at 25°C in a dark growth chamber. The infected calli were moved to a sterile plastic bottle and washed five times with sterile water to remove excessive *Agrobacterium*. After being dried on a sterilized filter paper, these calli were transferred onto screening medium at 28°C in a dark growth chamber for 5 weeks. During the screening stage, infected calli were transferred to fresh screening medium every two weeks. After the screening stage, actively growing calli were moved onto regenerative medium for regeneration at 28°C with a 16h light/8h dark cycle. After 3-4 weeks, transgenic seedlings were transferred to sterile plastic containers containing fresh solid medium and grown for 2-3 weeks before being transferred into soil. Transgenic rice plants were grown in a growth chamber at 28°C with a 16h light/8h dark cycle.

### Mutagenesis analysis at target sites

Genomic DNA was extracted from transgenic plants using the CTAB method^55^. The genomic region flanking the CRISPR target site for each gene was amplified and sequenced. Samples with heterozygous and biallelic mutations were decoded using CRISP-ID^56^.

### Whole genome sequencing and Data analysis

For each sample, about 1 g of fresh leaves were collected from seedlings between five and six weeks old. DNA samples were extracted using the Plant Genome DNA Kit (Tiangen) as described by the manufacturer. All 69 samples were sequenced by Bionova (Beijing, China) using the Illumina X10 platform. Adapters were trimmed using SKEWER (v. 0.2.2)^57^ and the Illumina TrusSeq adapter. Cleaned reads were mapped to rice reference sequence TIGR7 (http://rice.plantbiology.msu.edu/)^58^ with BWA (v. 0.7.15) software^59^. The Genome Analysis Toolkit (GATK)^60^ was used to realign reads near indels and recalibrate base quality scores by following GATK best practices^61^. A known SNPs and indels database for GATK best practices was downloaded from Rice SNP-Seek Database (http://snp-seek.irri.org/)^62^. Whole genome SNVs were detected by LoFreq^63^, MuTect2^64^ and VarScan2^65^. Whole genome indels were identified using MuTect2^64^, VarScan2 and Pindel^66^. Bedtools^67^ and BCFtools^68^ were used to process overlapping SNVs/indels. Off-target sites were predicted with CRISPOR^30^ online and Cas-OFFinder software^31^ by allowing up to 10 nt mismatch. Genome-wide map of mutations was plotted with Circos software^69^. Structural variants and translocation events were analyzed by using TopHat2^70^ with ‘—fusion-search’ parameter, DELLY^71^ with default parameter and manually checking with IGV software^72^. The NCBI BLAST+ with ‘-task blastn-short’ parameter was used for off-target mutations site analysis, which include mismatch, deletion and insertion. Reads mapping screenshots were from Golden Helix GenomeBrowse ^®^ visualization tool v2.1. Data processing and analyses were completed using R and Python. One T1 sample of Cas9-CC2 was excluded from analyses due to contamination of fungal DNA.

## ACKNOWLEDGEMENTS

This work was supported by the National Science Foundation of China (31330017 and 31771486) and the Sichuan Youth Science and Technology Foundation (2017JQ0005) to Y.Z. This work was also supported by grants including the National Transgenic Major Project (2018ZX08022001-003) to Y.Z., and T.Z., the Jiangsu Specially-Appointed Professor and the Priority Academic Program Development of Jiangsu Higher Education Institutions (PAPD) to T.Z., and startup funds provided by University of Maryland to Y.Q.

## AUTHOR CONTRIBUTIONS

Y.Z. proposed the project. Y.Q., T.Z., and Y.Z. designed the experiments. X.T., J.Z., D.Z., and A.M. generated CRISPR-Cas nuclease Vectors. X.T., J.Z., L.T., X.X., D.Z., and X.Z. generated stable transgenic rice and identified the rice mutants. X.T., J.Z., Q.R., Z.Z., B.L., and X.Z. prepared samples for WGS and confirmed WGS results with Sanger Y.AA. sequencing. G.L., Q.Y., Z.G., and T.Z. performed WGS data analysis. Y.Q., T.Z., and Y.Z. analyzed the data and wrote the manuscript. All authors participated in discussion and revision of the manuscript.

## SUPPLEMENTARY FIGURE LEGENDS

**Supplementary Fig. 1 Non-mosaic mutations at target sites revealed by Sanger sequencing in all Cas9 edited T0 lines chosen for WGS**

**Supplementary Fig. 2 Non-mosaic mutations at target sites revealed by Sanger sequencing in all Cpf1 edited T0 lines chosen for WGS**

**Supplementary Fig. 3 Venn diagram of SNVs and indels detected by multiple variant callers in WT plants for three consecutive generations**

**Supplementary Fig. 4 Venn diagram of SNVs and indels detected by multiple variant callers in four types of tissue culture related control samples**

**Supplementary Fig. 5 Genome-wide distribution of mutations from tissue culture only, Agro-infected, Cas9 backbone, Cpf1 backbone, Cas9 T0 and Cpf1 T0 samples.**

**Supplementary Fig. 6 Venn diagram of SNVs and indels detected by multiple variant callers in Cas9-edited T0 lines**

**Supplementary Fig. 7 Venn diagram of SNVs and indels detected by multiple variant callers in Cpf1-edited T0 lines**

**Supplementary Fig. 8 Mapped T-DNA insertion site for each T0 line.** T-DNA insertion sites are shown in black lines. Samples with more than 1 T-DNA insertion are marked in red.

**Supplementary Fig. 9 Alignment of top candidate off-target sites to the target sites based on WGS-discovered mutations in Cas9 T0 lines.** The putative off-target sites are aligned with the on-target sequences. They are identified based on meeting three criteria: (1) contain a PAM (NGG for Cas9), (2) allow up to 10 nt mismatches with the on-target sequences (20 nt for Cas9 and 23 nt for Cpf1) and (3) contain mutations in these sequences.

**Supplementary Fig. 10 Alignment of top candidate off-target sites to the target sites based on WGS-discovered mutations in Cpf1 T0 lines.** The putative off-target sites are aligned with the on-target sequences. They are identified based on meeting three criteria: (1) contain a PAM (TTTN for Cpf1), (2) allow up to 10 nt mismatches with the on-target sequences (20 nt for Cas9 and 23 nt for Cpf1) and (3) contain mutations in these sequences.

**Supplementary Fig. 11 Examples of mutation sites snapshot from Genome Browser**

**Supplementary Fig. 12 Germline transmitted mutations at target sites revealed by Sanger sequencing in all Cas9 edited T1 lines**

**Supplementary Fig. 13 Germline transmitted mutations at target sites revealed by Sanger sequencing in all Cpf1 edited T1 lines**

**Supplementary Fig. 14 Venn diagram of SNVs and indels detected by multiple variant callers in Cas9 T1 lines**

**Supplementary Fig. 15 Venn diagram of SNVs and indels detected by multiple variant callers in Cpf1 T1 lines**

**Supplementary Fig. 16 Analysis of mutations found in Cas9 and Cpf1 T0 and T1 lines.** Occurrence of SNV and Indel counts are calculated per 1 million base pairs (Mb) with five genome annotation units (Gene, TE, Repeats, CDS and Introns) for six different sample types. Note for the mutations identified in T1 plants were broken into two groups: those inherited from T0 plants and new mutations generated *de novo*. The average numbers of SNVs and indels of biological replicates are shown. Error bars indicate s.e.m.

**Supplementary Fig. 17 Genome-wide distribution of spontaneous (new) mutations discovered in WT across two generations as well as in Cas9 and Cpf1 T1 lines.**

**Supplementary Fig. 18 SNV rates and types in different treatments or generations.** Mutation rates of detected SNVs of all T0 and T1 plants. Control includes four control sample types: tissue culture only, Agro transformation, Cas9 backbone and Cpf1 backbone. Complementary mutations, such as A>C and T>G, are pooled.

## SUPPLEMENTARY TABLES

**Supplementary Table 1. Summary of all samples and target-site genotyping results**

**Supplementary Table 2. Summary and statistics of whole genome sequencing**

**Supplementary Table 3. Empirically validated off-target sites vs Cas-OFFinder or CRISPOR-predicted off-target sites**

**Supplementary Table 4. Validated mutations by Sanger sequencing Supplementary Table 5. List of other oligos used in this study**

